# The *Drosophila* wing is a high-throughput and versatile screening tool for Tau-mediated disease mechanisms and drug discovery

**DOI:** 10.1101/2025.05.21.655273

**Authors:** Miguel Ramirez-Moreno, Amber S. Cooper, Tianshun Lian, Jie Liu, Seyedehleila Abtahi, Efthimios M.C. Skoulakis, Lovesha Sivanantharajah, Douglas Allan, Amritpal Mudher

**Affiliations:** School of Biological Sciences, University of Southampton, Southampton, SO17 1BJ, UK; Department of Cellular and Physiological Sciences, Life Science Institute, University of British Columbia, Vancouver, V6T 1Z3, Canada; Institute for Fundamental Biomedical Research, Biomedical Sciences Research Centre “Alexander Fleming”, Vari, 16672, Greece; School of Biological Sciences, Bangor University, Bangor, LL57 2UW, UK

## Abstract

Tau protein contributes to microtubule stability, which is disrupted in Alzheimer’s disease and other Tauopathies. In these diseases, Tau molecules become hyperphosphorylated, misfolded and aggregated, propagating pathology across the brain. Studies dissecting disease mechanisms or screening disease-modifying therapies rely on animal models that unveil pathogenic events *in vivo* but also take several weeks or months to complete. Here we describe a versatile experimental paradigm that yields results in days and yet offers all the advantages of a genetically tractable *in vivo* system: the *Drosophila* wing disc. Mimicking neurotoxicity, human Tau expression causes cell death in the wing disc leading to quantifiable phenotypes in the adult wing. The neuroprotective peptide NAP ameliorates Tau toxicity in this system, validating it as a cost-effective drug screening tool. Phenocopying adult neurons, Tau toxicity in the wing disc is exacerbated by simulating hyper-phosphorylation and prevented by suppressing aggregation. Additionally, we show that the wing disc can dissect disease mechanisms that underpin clinically relevant Tau variants. Thus, the wing disc offers an *in vivo* experimental paradigm for fast and efficient exploration of disease mechanism and screening.

## Introduction

The developing wing of the fruit fly, *Drosophila* melanogaster, is a well-established paradigm for growth, morphogenesis, regeneration and tumorigenesis (Tripathi and Irvine, 2022, Beira and Paro, 2016). The adult organ develops from the larval imaginal wing disc, an epithelial sac that like the nervous system is originated from the embryonic ectoderm. During metamorphosis, the two wing discs give rise to both part of the thorax wall (notum region), the wing hinge and the actual wing blade (Beira and Paro, 2016, Tripathi and Irvine, 2022). The wing discs are relatively flat structures more easily extractable than brains and can be cultured *ex vivo* for short periods (Moreno et al., 2022, Handke et al., 2014). Using established genetic tools (like the Gal4/UAS system (Brand and Perrimon, 1993)), fly researchers can express genes of interest, and their ensuing protein products, within finite anatomically defined regions of the wing disc that can be traced to distinct regions of the adult wing (Tripathi and Irvine, 2022). Tractable genetic lineages divide the organ into compartments (anterior/posterior or dorsal/ventral), so researchers can target expression of genes of interest to specific compartments of the organ whilst using the other compartments as an internal experimental control within the same organism (Tabata et al., 1995, Brand and Perrimon, 1993). This highly accessible system lends itself to all manner of studies relevant to disease modelling, from dissecting disease mechanisms to candidate drug screening assays, all of which can be done in a week. This offers a great first-pass model system to streamline the design of long-term assays in more clinical-relatable paradigms like the adult nervous system, which require significantly longer experimental times.

Dementia is currently one of the most pressing health challenges worldwide, with the most common cause being Alzheimer’s Disease (AD). AD pathological hallmarks include the aberrant accumulation and dysfunction of the microtubule associated protein (MAP) Tau (Weingarten et al., 1975, Alzheimer, 1907, Parra Bravo et al., 2024). Under normal physiological conditions, Tau regulates the neuronal cytoskeleton, but in disease states, it accumulates to form aggregates like Neurofibrillary Tangles (NFTs) within the brain (Tabeshmehr and Eftekharpour, 2023, Goedert et al., 2023, Drechsel et al., 1992, Kanaan, 2023). Other neurodegenerative diseases characterized by Tau dysfunction, such as frontotemporal lobal degeneration, progressive supranuclear palsy or Pick’s disease are collectively termed as tauopathies (Zempel et al., 2017). A series of extrinsic and intrinsic factors, from the cellular environment to the Tau protein itself, contribute to its transition from a physiological to pathological state (Morris et al., 2011, Kanaan, 2023, Goedert and Jakes, 2005). Pathological features of Tau include abnormal phosphorylation (hyperphosphorylation), mislocalization, conformational changes and aggregation, occurring in a cascade of events leading to NFTs (Buchholz and Zempel, 2024a, Alonso et al., 2001, Cowan et al., 2010, Stoothoff and Johnson, 2005, Kimura et al., 2018, Wegmann et al., 2021). It is important to investigate why Tau becomes pathological and how it impacts affected cells, and to address this question many successful experimental paradigms of neurodegeneration have been established. However, most of these models examine Tau pathology in ageing nervous systems which often require significant time and resources—sometimes taking several years as with rodent models. Even short-lived invertebrate models, like *Drosophila* (Cowan et al., 2011, Sivanantharajah et al., 2019), have limitations for large scale screenings as neurodegeneration requires weeks to manifest. Whilst 2- or 3-dimensional cell culture models can be faster, they are *in vitro* experimental paradigms that fail to fully reproduce the complexity of animal brains and are prone to other technical issues such as inefficient transfection that could affect the reproducibility of assays.

Thus, we need a fast and versatile *in vivo* platform that combines the ability to dissect cellular pathways with a superior and efficient spatiotemporal genetic tractability. In this study, we validate the developing wing of *Drosophila* as a new tool that meets all these criteria. This experimental paradigm takes advantage of the extensive genetic toolkit of *Drosophila* and, as described in this work, can provide a fast, highly efficient and relevant system for the interrogation of Tau-mediated disease mechanisms and drug screening. In this system, the expression of human Tau isoforms triggers an apoptotic response in the wing disc, which correlates with the severity of observed phenotypes in the adult organism. We recapitulated several aspects of Tau-mediated toxicity that have been described in the adult central nervous system, highlighting the potential of the wing disc experimental paradigm to study impacts of Tau on cellular pathways. The extensive use of the wing disc to study cell biology (Moreno et al., 2022), and especially intracellular trafficking processes implicated in Tau toxicity (Yan and Zheng, 2021, Rauch et al., 2020), represents a further advantage of this platform. Additionally, we show the capacity of this model to rescue Tau phenotypes using drugs that have been shown to be effective against Tau-mediated neurotoxicity in rodent models, validating the use of the *Drosophila* wing for drug discovery and screening.

## Results

### The wing disc and adult wing are a fast and robust scorable paradigm to quantify Tau toxicity

In early embryogenesis, cells and tissues of the developing fly acquire identities that separate them into developmental compartments, delimited by boundaries of cells and lineages (Garcia-Bellido et al., 1973, Tabata et al., 1995). The GAL4/UAS system allows to specifically target, for example, the posterior compartment, constituted by *engrailed*-expressing cells (*en*-Gal4), leaving the anterior compartment as internal control (Fig. 1A)(Tabata et al., 1995, Brand and Perrimon, 1993). A change in the relative contribution of the posterior compartment to the total disc size (as measured with the posterior/anterior ratio, see Methods) allows the detection of growth/developmental anomalies at the targeted tissue (Moreno et al., 2022), while accounting for differences in absolute size due to diet or genetic background. We expressed mCherry-tagged (mCh::) copies of two 4R isoforms of human Tau (henceforth hTau): hTau0N4R and hTau2N4R (Corsi et al., 2022, Mandelkow and Mandelkow, 2012, Parra Bravo et al., 2024), which were inserted into the same chromosomal site (AttP40) to minimize background differences. The expression of hTau isoforms were compared to tissues with expression of mCherry protein alone, which caused no known deleterious effects (Fig. 1B)(Moreno et al., 2022). Both hTau isoforms led to a morphological change in the wing disc, reducing the size of the posterior compartment (Fig. 1B, C). Unexpectedly, hTau0N4R caused a stronger effect than the full-length hTau2N4R isoform (Fig. 1C). Flies expressing hTau at their posterior compartments reached adulthood with no developmental delay or detectable anatomical defects elsewhere, but those expressing hTau0N4R displayed a marked reduction of the posterior region of the adult wing, identified as the area below the vein L4 (Fig. 1A, D, E) Alongside the reduced size, many of the hTau0N4R-expressing compartments displayed notched margins (Fig. 1D). In comparison, the milder wing disc phenotype on hTau2N4R-expressing individuals was compensated by the organ and did not manifest after metamorphosis (Fig. 1D, E), indicating that the baseline insult due to full-length hTau2N4R expression can be compensated by the wing disc tissue.

**Fig. 1.**
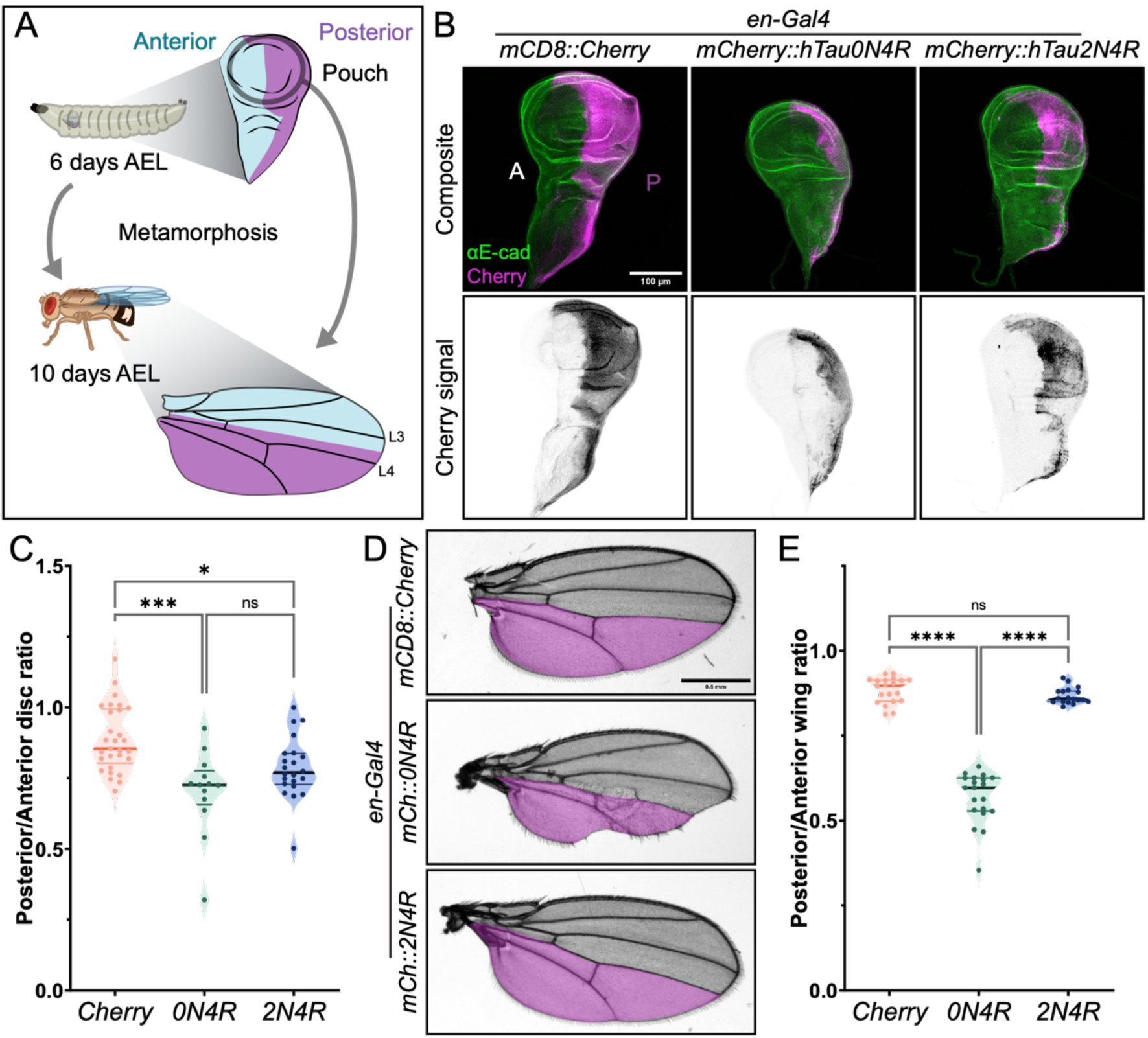
hTau expression reduces the compartment size at the Drosophila wing. (A) Cartoon depicting the developing third instar larvae (6 days after egg laying, AEL) and *Drosophila* adult organism, with views of the developing imaginal wing disc (top) and adult wing (bottom). Cells corresponding to the posterior compartment are highlighted in magenta, and the circle highlight the disc pouch, from where the adult wing blade originates during metamorphosis. (B) Wing discs expressing mCD8::Cherry (left) or mCherry-tagged (mCh::) hTau isoforms, showing mCherry (magenta, top; grayscale, bottom) and E-cadherin (E-cad) antibody (green, top) signal, with letter indicating anterior (A) and posterior (P) compartments. Scale bar: 100 µm. (C) Posterior/Anterior ratios of control (mCD8::Cherry) and mCh::hTau-expressing wing discs. Dots represent individual discs (n= 29, 15 and 22). *p<0.05 and ***p<0.001 (Kruskal-Wallis test). Both 0N4R and 2N4R isoforms significantly reduce compartment size when compared to the control. (D) Adult wings (grayscale) expressing mCD8::Cherry (top) or mCh::hTau isoforms, showing a magenta overlay of the approximated posterior compartment (below L4). Scale bar: 0.5 mm. (E) Posterior/Anterior ratios of adult wings expressing mCD8::Cherry or mCh::hTau isoforms during development. Dots represent individual wings (n= 20, 20 and 18). ****p<0.001 (Kruskal-Wallis test).

With such effects at the adult wing blade, we analysed the death rate at the wing disc pouch, the region it arises from during metamorphosis. To assess the amount of apoptosis at the wing disc, occasional but sparse during normal development, we analysed the signal of cleaved effector caspase Dcp-1, one of the Caspase-3-like proteins of *Drosophila* and downstream actor of apoptosis (Fig. 2A) (Song et al., 1997), at the basal region of the discs where dying cells are extruded from the tissue (Moreno et al., 2022, Bergantiños et al., 2010, Matamoro-Vidal et al., 2024) (Fig. 2B). The pouch region of each compartment was identified using anatomical and fluorescence markers delineating its boundaries (Fig. 2C). While the expression of mCherry alone did not trigger apoptosis, expression of both hTau isoforms led to significant apoptosis in the posterior compartment (Fig. 2C, D). The stronger phenotype upon hTau0N4R expression was consistent with the more pronounced tissue-level phenotypes at larval and adult stages (Fig. 1), thus correlating the reduced compartment size with apoptosis activation and cell death.

**Fig. 2.**
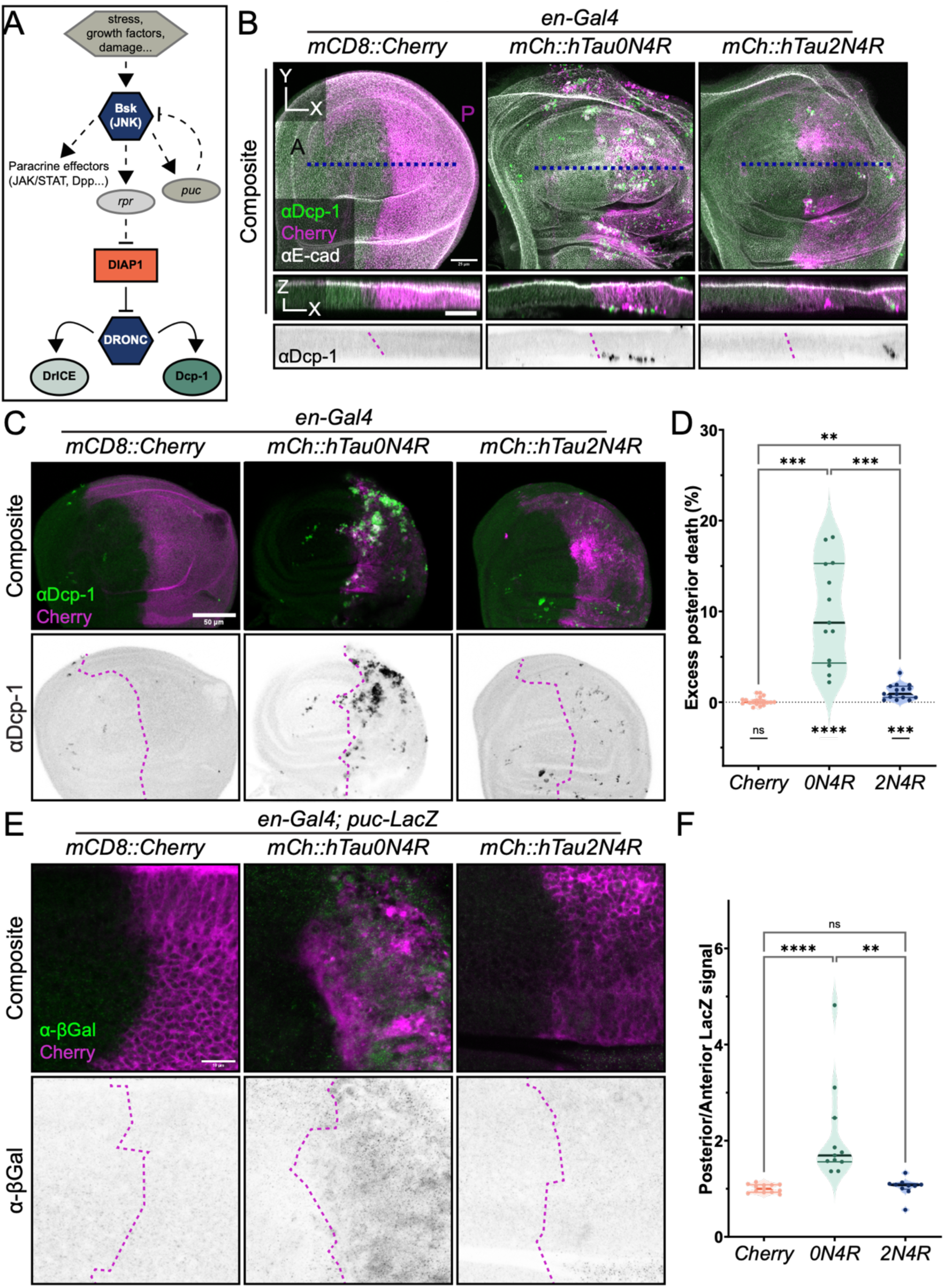
hTau expression induces apoptotic response at the wing disc. (A) Simplified scheme of the *Drosophila* apoptotic pathway. Briefly, pro-apoptotic factors activate among others JNK-signaling (Basket, BSK, as the Drosophila homologue) which in turn promotes the expression of activator caspases such as Reaper (*rpr*) and the expression of the negative regulator Puckered (*puc*). Initiator caspases inhibit *Drosophila* Inhibitor of Apoptosis 1 (DIAP1), which respectively inactivates Dronc (*Drosophila* homologue of Caspase9). Dronc then cleaves Caspase-3 like proteins such as Dcp-1 and Drice. (B) Wing pouches expressing mCD8::Cherry (left) or mCherry-tagged (mCh::) hTau isoforms, showing cleaved Dcp-1 antibody (green, top and middle; grayscale, bottom sagittal projection), mCherry (magenta, top and middle) expressed at the posterior compartment (P) and E-cadherin antibody (white, top and middle) signal. Sagittal views are the bottom represent a projection of 3.78 µm at the Y position of the blue dashed line in top panel. Scale bars: 25 µm. (C) Wing pouches expressing mCD8::Cherry (left) or mCherry-tagged (mCh::) hTau isoforms, showing cleaved Dcp-1 antibody (green, top; grayscale, bottom) and mCherry (magenta, top) signal. Magenta dashed lines at the bottom panel indicate the Anterior-Posterior compartment boundary. Scale bar: 50 µm. (D) Excess of apoptotic area at the posterior compartment for the indicated genotypes. Dots represent individual discs (n= 19, 13 and 15). *p<0.05, **p<0.01, ***p<0.001 and ****p<0.0001 (Brown-Forsythe and Welch ANOVA test for multiple comparisons, and one-sample *t* test in comparison to zero for random distribution of apoptosis). (E) Apical region of dorsal wing pouch cells expressing mCD8::Cherry (left) or mCherry-tagged (mCh::) hTau isoforms alongside the *puc*-lacZ sensor, showing β-galactosidase antibody (β-gal, green, top; grayscale, bottom) and mCherry (magenta, top) signal. Magenta dashed lines at the bottom panels indicate the Anterior-Posterior compartment boundary. Scale bar: 10 µm. (F) Posterior/Anterior ratio of β-galactosidase antibody in dorsal wing pouch cells. Dots represent individual discs (n= 13, 11 and 10). **p<0.01 and ****p<0.0001 (Kruskal-Wallis test).

JNK-signalling is responsible for the release of multiple signals in the developing wing disc, including apoptosis and compensatory proliferation following stress (Pérez-Garijo et al., 2009, Moreno et al., 2022, Pinal et al., 2019). JNK-signalling can be tracked with the expression of *puckered* (*puc*), which encodes a negative regulator that acts as negative feedback upon pathway activation (Fig. 2A) (Martín-Blanco et al., 1998, Ring and Martinez Arias, 1993). We used the *pucE69-lacZ* construct to drive expression of β-galactosidase (Ring and Martinez Arias, 1993) and found that overexpression of hTau0N4R, but not hTau2N4R, was sufficient to cause tissue-wide upregulation of JNK-signalling (Fig. 2E, F). Overall, we found that the expression of hTau induces toxicity in the developing wing disc, which is akin to Tau-induced neurotoxicity and can be quantified with multiple readouts in both the larval discs and adult wing.

### The Drosophila wing can be used for testing drugs counteracting Tau toxicity

As the generation of disease-modifying treatments represents one of the biggest challenges faced by dementia researchers and having a high-throughput drug screening platform is highly desirable, we asked whether the Tau-induced cytotoxicity at the wing disc could be ameliorated using drug treatments. Previous studies have highlighted the neuroprotective properties of the octapeptide NAPVSIPQ (NAP, Davunetide), an active component of Activity-Dependent Neuroprotective Protein (ADNP), essential for mammalian brain development (Bassan et al., 1999, Gozes et al., 2005). Indeed, we previously reported that administration of NAP on the diet was sufficient to improve phenotypes associated with hTau expression in the nervous system, including defects in larval locomotion and axonal transport (Quraishe et al., 2016, Quraishe et al., 2013). To test whether diet-administered NAP would reach the imaginal tissue as well, we raised flies expressing hTau0N4R at their posterior compartments with the peptide dissolved in the food, once we had confirmed that the highest dosage of 25 µg/ml was harmless to the control discs (Fig. S1). There was an improvement of the tissue-level phenotype with the smallest concentration, 5 µg/ml, but not with the high dosage (Fig. 3A, B). These results correlate with the apoptotic rates under these conditions (Fig. 3C, D). NAP administration improved the morphology of the adult wing again in the small concentration sample (Fig. 3E, F), while the high dose caused a striking bi-modal distribution of adult wings, with some of them displaying almost *wild type*-like wings. Therefore, the improvement of the phenotypes caused by hTau0N4R expression highlight the potential of the wing disc as a drug screening tool for finding mediators and inhibitors of Tau-derived toxicity.

**Fig. 3.**
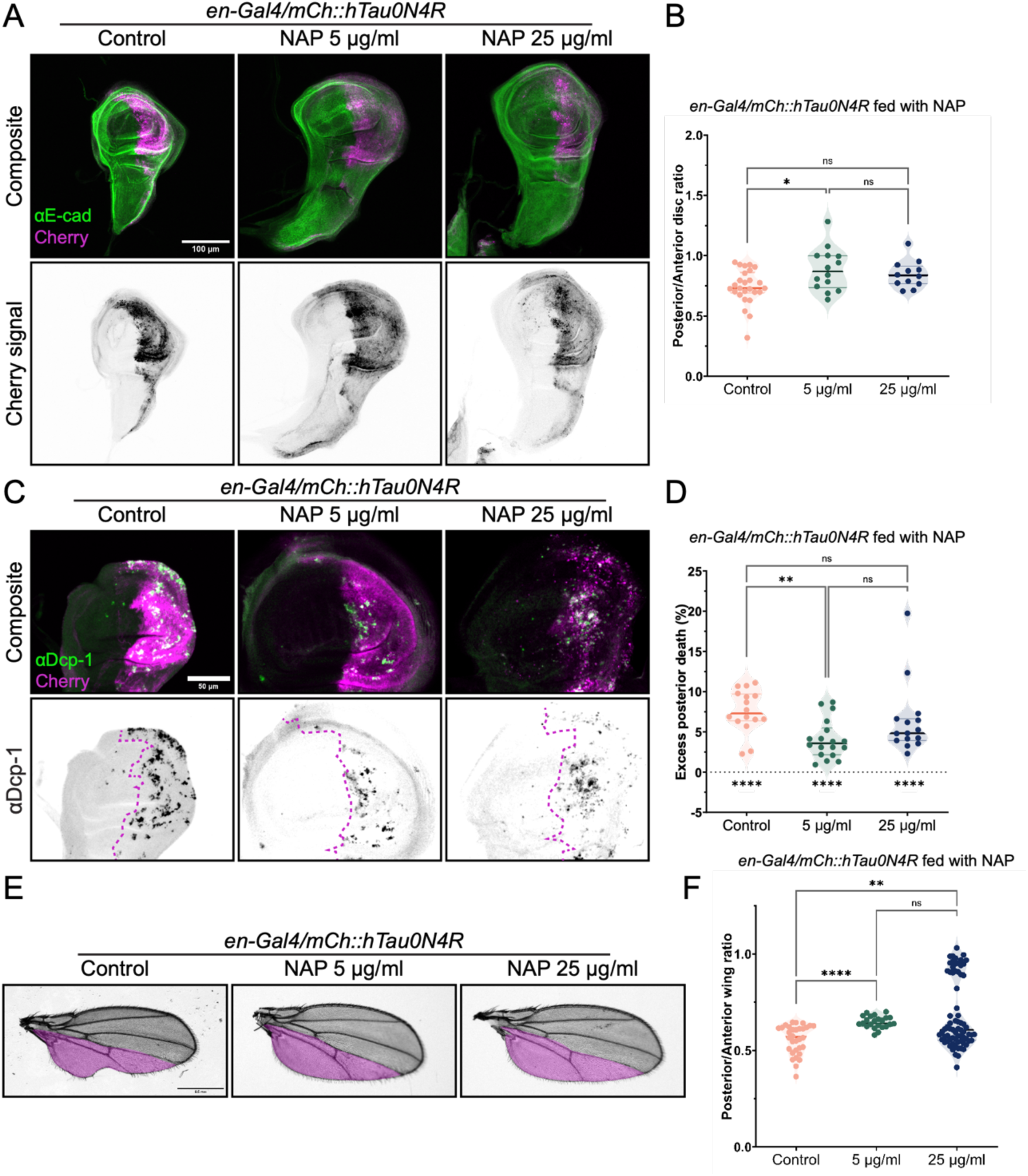
NAP peptide partially rescues hTau toxicity at the Drosophila wing. (A) Wing discs expressing mCherry-tagged (mCh::) hTau0N4R, corresponding to individual treated with NAP peptide, showing Cherry signal (magenta, top; grayscale, bottom) and E-cadherin (E-cad) staining (green, top). Scale bar: 100 µm. (B) Posterior/Anterior ratios of NAP-treated mCh::hTau0N4R-expressing wing discs. Dots represent individual discs (n= 16, 17 and 15). *p<0.05 (Kruskal-Wallis test). (C) Wing pouches expressing mCh::hTau0N4R, corresponding to individual treated with NAP peptide, showing cleaved Dcp-1 antibody (green, top; grayscale, bottom) and mCherry (magenta, top). Magenta dashed lines at the bottom panel indicate the Anterior-Posterior compartment boundary. Scale bar: 50 µm. (D) Excess of apoptotic area at the posterior compartment for the indicated genotypes. Dots represent individual discs (n= 16, 18 and 15). **p<0.01 and ****p<0.0001 (Kruskal-Wallis test for multiple comparisons, and one-sample *t* test in comparison to zero for random distribution of apoptosis). (E) Adult wings (grayscale) of NAP-treated flies expressing mCh::hTau0N4R during development, showing a magenta overlay of the approximated posterior compartment (below L4). Scale bar: 0.5 mm. (F) Posterior/Anterior ratios of NAP-treated flies expressing mCh::hTau0N4R. Dots represent individual wings (n= 32, 23 and 72). **p<0.01 and ****p<0.001 (Kruskal-Wallis test).

### The Drosophila wing can be used to genetically dissect novel mechanisms of Tau toxicity

So far, we have characterized the baseline toxic response caused by hTau expression at the developing wing, and its sensitivity to drug treatments. We next asked whether pathogenic features of hTau such as aggregation or hyper-phosphorylation, could be targeted to modify its toxicity in this system. Regarding aggregation, two hexapeptide motifs (VQ domains) inside the microtubule-binding domain (MTBD), ^275^VQIINK^280^ and ^306^VQIVYK^311^ are of special interest as they comprise the core of the pathological Tau fibrils (Wu et al., 2022, von Bergen et al., 2000). Recently, we showed that the deletion of the ^306^VQIVYK^311^ aggregation motif was sufficient to render human Tau inert in the ageing *Drosophila* nervous system (Cooper et al., 2024).

We sought to test whether phosphorylation would also increase toxicity at the wing disc, and if the capacity for self-aggregation was a contributor to toxicity in this system as well. We first tested the effects of deleting the ^306^VQIVYK^311^ domain (ΔVQIVYK), one of the two aggregation promoting motifs shown to have the highest propensity for aggregation *in vitro* (Ganguly et al., 2015). hTau2N4R^ΔVQIVYK^ completely rescued the phenotype on both cell and tissue levels at the wing disc, effectively eliminating the Tau-mediated toxicity (Fig. 4A-D). We next independently deleted the other aggregation motif, ^275^VQIINK^280^ (ΔVQIINK), an hexapeptide with stronger intrinsic self-aggregation capacity but more dispensable than VQIVYK in the Tau pathological cascade (Seidler et al., 2018, Passarella and Goedert, 2018). hTau2N4R^ΔVQIINK^ led to a similar rescue of the basal hTau toxicity (Fig. 4A-D). To obtain more information regarding the mechanisms of hTau-induced toxicity, we simulated hyperphosphorylation with the E14 variant, which carries 14 serine/threonine to glutamate substitutions in residues linked to pathology, resulting in a phosphomimic protein with increased neurotoxicity in multiple models (Hatch et al., 2017, Steinhilb et al., 2007, Cooper et al., 2024, Hoover et al., 2010). hTau2N4R^E14^ did not ostensibly impact the size of the developing posterior compartment (Fig. 4A, C), but triggered a strong apoptotic response in the posterior pouch (Fig. 4B, D). We then assessed the impact of independently deleting the two aggregation motifs on the E14 wing phenotypes, following our discovery that hTau2N4R^E14.ΔVQIVYK^ is an inert protein unable to drive neurodegeneration in the fly (Cooper et al., 2024). We recapitulated the total rescue of the E14 variant in the developing wing with hTau2N4R^E14.ΔVQIVYK^, but strikingly, hTau2N4R^E14.ΔVQIINK^ failed to completely abrogate the cytotoxic response of the E14 variant (Fig. 4A-D). The results of these variants at the adult wing reproduced those in the wing discs, with hTau2N4R^E14^ manifesting a marked phenotype at the targeted compartment that was, again, rescued by the two deletions but only completely by ΔVQIVYK (Fig. 4E, F).

**Fig. 4.**
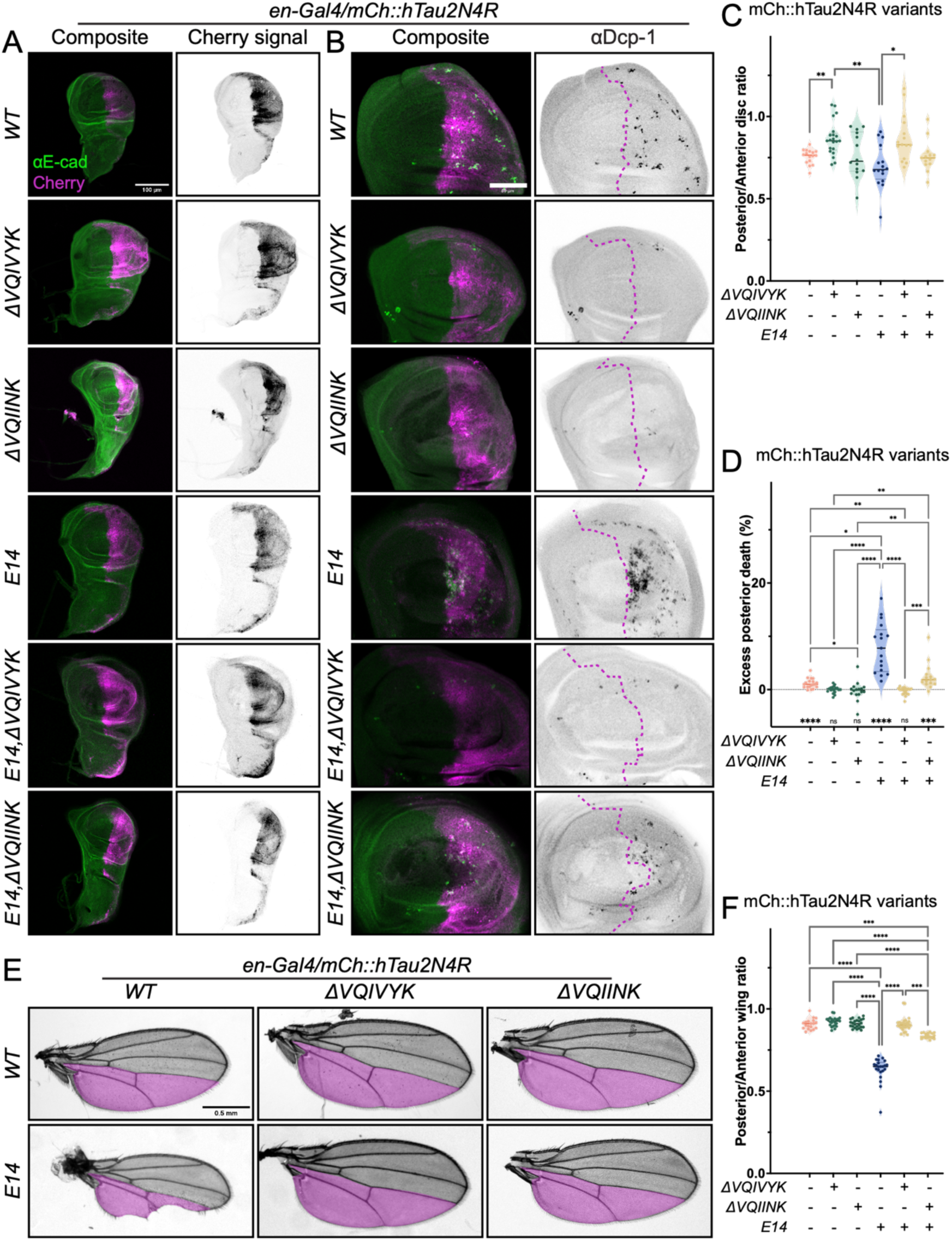
hTau requires the VQIVYK motif to drive toxicity at the developing wing. (A) Wing discs expressing variants (rows) of the original mCherry-tagged (mCh::) hTau2N4R isoform (WT, top), showing Cherry signal (magenta, left; grayscale, right) and E-cadherin (E-cad) staining (green, left). Scale bar: 100 µm. (B) Wing pouches expressing mCh::hTau2N4R variants, showing cleaved Dcp-1 antibody (green, left; grayscale, right) and mCherry (magenta, left) signal. Magenta dashed lines at the right panel indicate the Anterior-Posterior compartment boundary. Scale bar: 50 µm. (C) Posterior/Anterior ratios of wing discs expressing mCh::hTau2N4R variants, showing design combinations at the bottom of the table. Dots represent individual discs (n= 16, 21, 14, 16, 15 and 13 discs). *p<0.05 and **p<0.01 (Brown-Forsythe and Welch ANOVA test). (D) Excess of apoptotic area at the posterior compartment for the indicated genotypes. Dots represent individual discs (n= 19, 20, 16, 17, 19 and 15). *p<0.05, **p<0.01, ***p<0.001 and ****p<0.0001 (Kruskal-Wallis, and one-sample Wilcoxon test comparison to zero for random distribution of apoptosis). (E) Adult wings (grayscale) expressing mCh::hTau2N4Rvariants (presence/deletion of aggregation motifs, rows; inclusion of phosphomimic E14 substitutions, columns), showing a magenta overlay of the approximated posterior compartment (below L4). Scale bar: 0.5 mm. (F) Posterior/Anterior ratios of adult wings expressing mCh::hTau2N4Rvariants during development. Dots represent individual wings (n= 19, 23, 24, 22, 28 and 21). ****p<0.001 (Kruskal-Wallis test).

We confirmed some of these results by introducing the sequence changes in the hTau0N4R isoform independently, which itself is more toxic than 2N4R (Fig. 1). We found that ΔVQIVYK in expressed hTau0N4R fully restored the wild type phenotypes at the developing wing disc, the apoptotic rate of the expressing cells and the adult wing structure (Fig. S2). In contrast, the E14 substitutions exacerbated the toxic response, causing a massive apoptotic response and a further reduction of the posterior compartment at the adult wing (Fig. S2).

Altogether, these assays demonstrate that the *Drosophila* wing is a valid paradigm to study disease mechanisms associated with Tau-mediated toxicity. We tested this hypothesis by recapitulating our results at the ageing nervous system (Cooper et al., 2024), confirming that the ^306^VQIVYK^311^ motif is essential for hTau to be toxic in this system as well, independently of the presence of pathology-prone phosphomimic substitutions. We also found that deletion of ^275^VQIINK^280^ failed to completely rescue the hypertoxic effects of the E14 substitutions, thus using our new platform to obtain new *in vivo* evidence of the differential contribution of both aggregation motifs to Tau toxicity.

### The wing disc can be used to study clinical variants of interest

We next sought to investigate whether our experimental paradigm could be used to study the toxicity of clinically relevant mutants and unpick the underlying mechanisms of their toxicity. We chose R406W, one of the many mutations linked to frontotemporal dementia with parkinsonism linked to chromosome 17 (FTDP-17), a dominantly inherited early onset form of dementia (Wittmann et al., 2001, Hutton et al., 1998, Goedert and Jakes, 2005).

In line with previous reports from *Drosophila* models expressing hTau2N4R^R406W^ in the aging nervous system, we found a stronger toxic response than to hTau2N4R (Fig. 5). This was abrogated by the additional substitution of S262 and S356, two targets of PAR-1 kinase, to alanine (hTau2N4R^S2A^) that renders them non-phosphorylatable, in the same protein (Fig. 5C, D). In the adult wings, examination of all the genotypes together indicated that expression of both S2A mutants (alone or alongside R406W) produced bigger posterior compartments than R406W alone (Fig. 5E, F), while no obvious defects on tissue size were found with R406W, as observed with the E14 variant (Fig. 4). Compared with discs expressing hTau2N4R, hTau2N4R^R406W^-expressing wings discs had increased apoptotic response at their posterior compartment (t=2.534, df=35, p=0.0159), and affected adult wings (t=2.260, df=29, p=0.0295). Importantly, we found that the S2A substitutions alone improved the basal hTau2N4R phenotype (Fig. 5) These results indicate that the two PAR-1 phosphorylation targets of hTau sequence are major contributors to Tau toxicity at the wing disc. Furthermore, as exemplified by R406W mutation, the wing disc allows for the discrimination of point mutations, even with simple readouts as the analysis of the adult wing using developmentally robust region cues such as the veins. In turn, the wide range of disease factors related to Tau protein that can be recapitulated at the developing wing of the fly certifies the promising usage of this system as a screening tool to aide *Drosophila* biologists and non-fly researchers alike in the discovery of disease mechanisms and windows to target them.

**Fig. 5.**
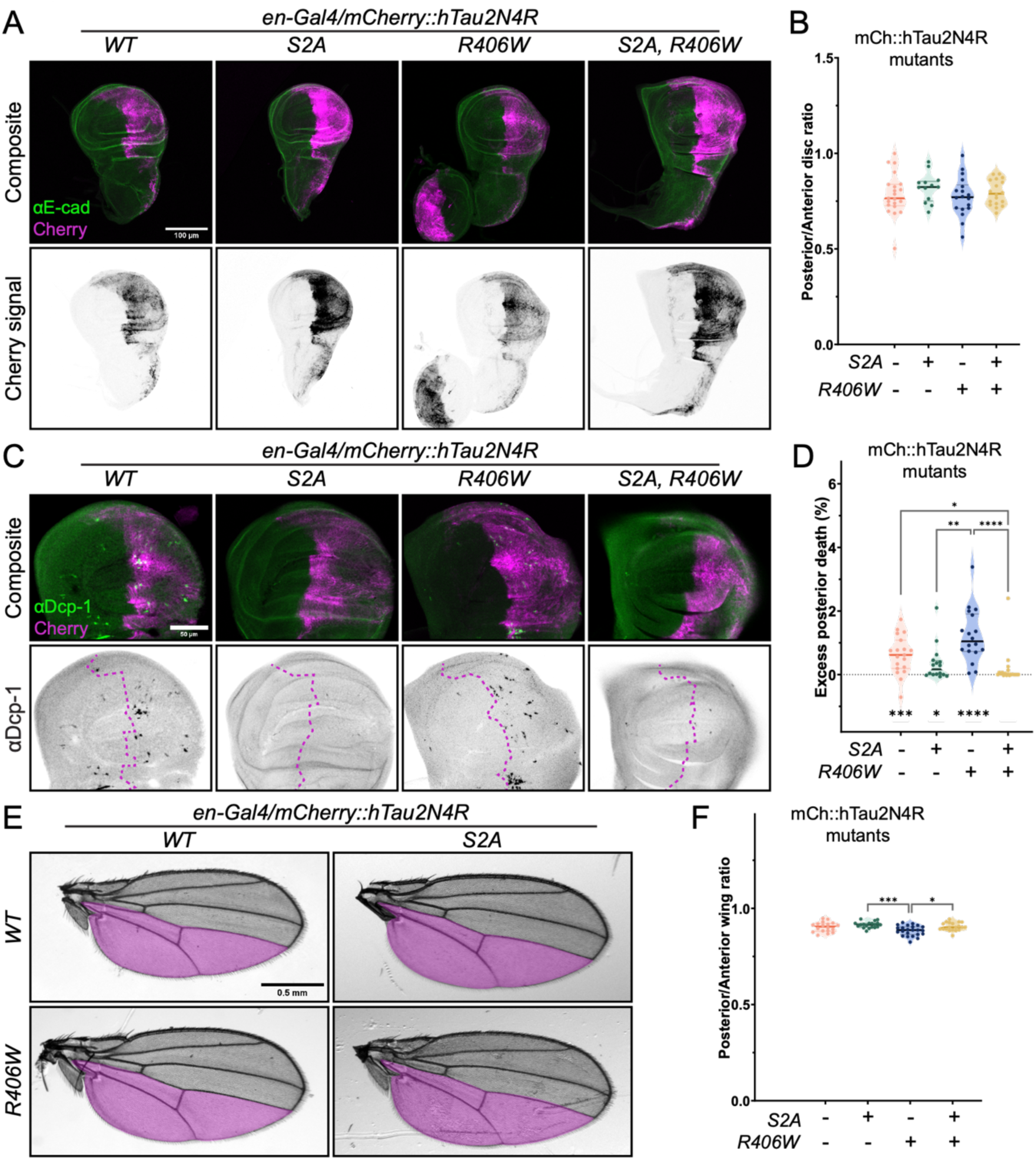
The wing disc allows the screening of point mutations in hTau sequence. (A) Wing discs expressing mutant variants (columns) of the original mCherry-tagged (mCh::) h Tau2N4R (WT, left), showing mCherry signal (magenta, top; grayscale, bottom) and E-cadherin (E-cad) staining (green, top). Scale bar: 100 µm. (B) Posterior/Anterior ratios of wing discs expressing mCh::hTau2N4R mutations, showing design combinations at the bottom of the table. Dots represent individual discs (n=21, 13, 18 and 18). (C) Wing pouches expressing mCh::hTau2N4R mutations, showing cleaved Dcp-1 antibody (green, top; grayscale, bottom) and mCherry (magenta, top) signal. Magenta dashed lines at the bottom panel indicate the Anterior-Posterior compartment boundary. Scale bar: 50 µm. (D) Excess of apoptotic area at the posterior compartment for the indicated genotypes. Dots represent individual discs (n= 19, 16, 18, 19). *p<0.05, **p<0.01, ***p<0.001 and ****p<0.0001 (Kruskal-Wallis test, and one-sample Wilcoxon test comparison to zero for random distribution of apoptosis). (E) Adult wings (grayscale) expressing mCh::hTau2N4R mutations, showing a magenta overlay of the approximated posterior compartment (below L4). Scale bar: 0.5 mm. (F) Posterior/Anterior ratios of adult wings expressing mCh::hTau2N4R during development. Dots represent individual wings (n= 19, 23, 22 and 28). *p<0.05 and ***p<0.001 (Brown-Forsythe and Welch ANOVA test).

## Discussion

The current work assessed the potential of the *Drosophila* wing to serve as a platform to dissect human Tau toxicity and the diverse mechanisms that underpin it. We show that in our system it is possible to recapitulate the cellular damage caused by hTau in fly and rodent models of neurodegeneration (Fig. 1 and 2). Importantly, clinically relevant mechanisms of toxicity—hyperphosphorylation and aggregation—identified in both invertebrate and vertebrate neurons also drive hTau toxicity in the *Drosophila* wing (Fig. 4). Furthermore, the administration of neuroprotective compounds that counteract Tau-mediated neurodegeneration in rodent models of Tauopathy suppressed hTau toxicity in the *Drosophila* wing, validating its utility as a drug discovery tool (Fig. 3). Finally, the wing can also be used to interrogate familial hTau mutations, underscoring its potential to study hTau variants of unknown clinical significance (Fig. 5). Taken together, we propose that the *Drosophila* wing is a powerful and accessible experimental paradigm yielding results consistent with those from neuronal models. Thus, it serves as a dependable and economical alternative to rapidly dissect disease mechanisms and perform high-throughput genetic/pharmacological screening.

### The wing disc is a robust tool to study Tau toxicity

Our study, focused on dissecting disease mechanisms underpinning Tauopathies, further highlights the usefulness of the *Drosophila* wing system in this area of research. Our results are consistent with reported effects attributed to hTau0N4R expression at the *patched* domain of the wing (Bougé and Parmentier, 2016). Others have additionally found that the wing cells are sensitive to the expression of Amyloid β42 and the dysregulation of its precursor protein (Fossgreen et al., 1998, Arnés et al., 2017). The wing has also been successfully employed to model disease mechanisms and identifying novel modifiers of C9ORF72 and TDP-43 (Lopez-Gonzalez et al., 2019, Yusuff et al., 2023).

Our observations suggest that ectopic cell apoptosis is the main consequence of hTau-mediated toxicity in the wing. Apoptosis in cells undergoing neurodegeneration has been linked to an attempt to re-enter the cell cycle (Frost, 2023, Cotman and Anderson, 1995, Troy and Jean, 2015, Khurana and Feany, 2007). Furthermore, Tau dysregulation has been linked to both promotion and inhibition of apoptosis (Li et al., 2007, Cimini et al., 2022). The most direct connection between Tau and apoptosis links DNA damage with the tumour suppressor p53 protein, which also controls the apoptotic response in *Drosophila* (Hooper et al., 2007, Sola et al., 2020, Farmer et al., 2020, Asada-Utsugi et al., 2022). While flies lack the negative p53 regulator MDM2 (which is functionally replaced by the *Drosophila* Corp), it is known that p53 promotes Tau phosphorylation levels, acting as a positive feedback for neurodegeneration (Jazvinšćak Jembrek et al., 2018, Hooper et al., 2007, Chakraborty et al., 2015).

Intriguingly, we found a difference in the intrinsic toxicity of 0N4R and 2N4R isoforms (Fig. 1, 4 and S2). Compared to the studies exploring the MTBD-related differences between 3 and 4R isoforms, the effects of the N repeats in the N-terminal region remain less explored, as many of the disease-relevant residues are common across all six isoforms (Cario et al., 2022, Kanaan et al., 2012). The different number of N repeats, resulting from alternative splicing affecting exons 2 and 3 (Goedert et al., 1989, Goedert and Jakes, 2005, Buchholz and Zempel, 2024a), has been linked to subcellular location differences (somatic versus axonal accumulation)(Zempel et al., 2017, Bachmann et al., 2021, Liu and Götz, 2013), affinity for binding partners (Liu et al., 2016, Buchholz and Zempel, 2024a) and opposing effects in promoting self-aggregation (Zhong et al., 2012). Although hTau0N4R and hTau2N4R display the same affinity for tubulin *in vitro* (Goode et al., 2000), the varying lengths of the N-terminal region may impact the ability of Tau to space microtubule bundles or cause crowding (Chung et al., 2016, Méphon-Gaspard et al., 2016, Prezel et al., 2018). Collectively, these differences in the physiological properties of the six hTau isoforms may be responsible for the isoform-specific pathogenic effects of Tau in neurons that have previously been reported by us and others (Vourkou et al., 2022, Buchholz et al., Zempel et al., 2017, Buchholz and Zempel, 2024b, Kosmidis et al., 2010).

Our analysis pipeline currently relies on simple image analysis using the stereotypical anatomy of an accessible tissue and the detection of cell death, as the apoptotic rate is minor in this tissue during physiological conditions except for specific small hotspots (Milán et al., 1997, Matamoro-Vidal et al., 2024). The availability of multiple Gal4 drivers that target other traceable compartments of the wing such as the anterior (*cubitus interruptus*) or dorsal (*apterous*) compartments, as well as the capacity to integrate different targets and expression of genes of interest thanks to the combination of multiple expression systems add further versatility to the platform (Tripathi and Irvine, 2022, Zirin et al., 2024). This repertoire of tools can be extended to interrogate multiple aspects of cell biology, for which this system has been widely used for decades, including endocytosis/intracellular trafficking (Moreno et al., 2022, Gao et al., 2017) and proteostasis (Joy et al., 2021), which are emerging areas of research in Tau pathology and its propagation (Yan and Zheng, 2021, Zhao et al., 2021, Papanikolopoulou and Skoulakis, 2020).

### Disease-relevant mechanisms mediate Tau toxicity in the wing

Alongside the experimental advantages, the key highlight of the wing system is that it recapitulates many features of hTau-related neurodegeneration. Much like in neuronal models, phosphorylation also plays a major role in promoting hTau-induced toxicity in the wing. This is exemplified with the behaviour of the phosphomimic E14 variant (Hoover et al., 2010), which exacerbates the toxicity of two different hTau isoforms (Fig. 4 and S2). It is also further supported with the suppression of toxicity attributed to the hypophosphorylated hTau2N4R^S2A^ variant (Fig. 5)(Chatterjee et al., 2009). On the other hand, and consistent with observations by us and others in the *Drosophila* neurons (Cooper et al., 2024, Passarella and Goedert, 2018), the presence of the VQIVYK aggregation motif is essential to drive toxicity for both hTau2N4R and hTau0N4R isoforms (Fig. 4 and S2). Indeed, we confirmed in our new system that the hexapeptide deletion can completely overcome the pathogenic effects of mimicking phosphorylation at 14 GSK3β sites in hTau2N4R, rendering this mutant (hTau2N4R^E14.ΔVQIVYK^) inert, exactly as is the case in the *Drosophil*a central nervous system (Cooper et al., 2024). The complete rescue observed with hTau0N4R^ΔVQIVYK^ is consistent with the finding that the C-terminal half of this isoform (which contains this domain) is required to drive toxicity in the wing margin tissue (Bougé and Parmentier, 2016). Interestingly, whereas the deletion of VQIVYK domain caused a 100% suppression of E14 toxicity, the deletion of the VQIINK aggregation-promoting domain, which is found exclusively in the 4R hTau isoforms, rescued approximately 80% of hTau2N4R^E14^-driven toxicity (Fig. 4)(Seidler et al., 2018). The capacity of the two VQ motifs to promote Tau aggregation differs depending on their molecular context *in vitro*, but both have been found to contribute to aggregation *in vivo* (Macdonald et al., 2019, Wu et al., 2022, Seidler et al., 2018, Ganguly et al., 2015). Adding to this growing literature, our data in the *Drosophila* wing system highlights the importance of targeting these two motifs as promising therapeutics to counteract Tau toxicity (Aggidis et al., 2024), and our platform as a fast and high-throughput platform to enable this.

The enhanced toxicity of the clinically relevant R406W mutant and its rescue upon the non-phosphorylatable S262 and S356 residues (Fig. 5) provides further evidence that the *Drosophila* wing can recapitulate the requirement of specific phospho-epitopes in Tau-mediated neurodegeneration (Parra Bravo et al., 2024, Chatterjee et al., 2009). Furthermore, these data show that the system can be used as an appropriate *in vivo* platform for understanding the pathogenic significance of multiple post-translational modifications of Tau protein that may mediate disease mechanisms (Min et al., 2015, Wada et al., 2024, Losev et al., 2021, Ait-Bouziad et al., 2020, Acosta et al., 2022).

### The Drosophila wing as a tool for drug discovery

Our findings that the neuroprotective peptide NAP (Leker et al., 2002, Oz et al., 2012, Oz et al., 2014, Quraishe et al., 2013) reduces the toxicity of hTau0N4R in our experimental paradigm (Fig. 3) demonstrates the utility of our system for screening Tau-targeting drugs. The easy observation of the highly stereotypical adult wing, which supports further anatomical analysis (Sonnenschein et al., 2015) lends itself to high-throughput screening of multiple drug panels. The advantages of *Drosophila* as an animal model have facilitated its employment in screenings related to cancer, inflammatory bowel disease and diabetes among others (La Marca et al., 2023, Richardson et al., 2015, Munnik et al., 2022, Xiu et al., 2022, Willoughby et al., 2013, Lagunas-Rangel et al., 2023). The wing model itself has been used as a successful platform for drug discovery and validation of EGFR inhibitors and modulators of metabolic diseases (Aritakula and Ramasamy, 2008, Bai et al., 2018, Merigliano et al., 2018)

### Conclusion

In conclusion, our study presents a novel platform to dissect mechanisms of disease underpinning neurodegeneration in Tauopathies. This highly accessible and versatile system can support Tau researchers in assessing factors that promote or reduce the pathological potential of the microtubule-associated protein, finding new interactors that mediate these effects, or exploring drug targets in a more efficient and affordable manner.

## Materials and methods

### Drosophila genetics and husbandry

*Drosophila melanogaster* flies were reared on standard media at 25°C. To express tools of interest, the GAL4/UAS system was employed (Brand and Perrimon, 1993) to drive expression at the posterior wing compartment using *engrailed*-GAL4. All hTau lines used (isoforms and variants) were tagged with N-terminal mCherry. New hTau transgenic lines were produced by the Allan Laboratory (University of British Columbia, Canada) by fusing hTau sequences downstream of 10XUAS sequences, and insertion into *attP40* (position 25C on chromosome II) to minimize position effect variegation. hTau isoform and variants cDNAs were subcloned using the BgIII site at the pCaU4B2G10XU plasmid using Gibson Assembly (NEB). Injection and establishment of stable transgenic lines were completed by GenomeProlab (Montreal, Canada). Precise transgenic lines and constructs are detailed in Resources Table.

### Drug assays

NAP (davunetide) (Peptide Protein Research, Fareham, UK) was added to fly food at concentrations of 5 μg ml–1 or 25 μg ml–1.

### Wing disc dissection and immunostaining

Third instar larvae (approximately 144 hours after egg laying at 25°C) were dissected in phosphate buffer saline (PBS). Cuticles with attached imaginal discs were fixed for 20 minutes with 4% paraformaldehyde (Electron Microscopy Sciences; 15713-S) in PBS at room temperature, then washed with PBS with 0.1% Triton X-100 (PBST, Merk; X100-1L). Cuticles were blocked for 45 minutes with 1% normal goat serum (NGS, Merk; G9023-10ML) in PBST at room temperature, and incubated with primary antibodies in PBST overnight at 4°C with constant agitation. The next day, cuticles were washed with PBST, incubated with secondary antibodies with 1% NGS in PBST for 2 hours at room temperature and washed in PBST, then transferred to PBS for wind disc extraction. Discs were transferred to the microscope slide and mounted in Prolong Glass (Invitrogen; P36980) and equilibrated for 72 hours in the dark at room temperature before imaging. All antibodies and concentrations used are listed in Resources Table.

### Adult wing imaging

Male adult flies were frozen upon collection up to 48 hours after hatching. Wings were removed and imaged using a Leica Stereoscopic microscope (LAS Software).

### Fluorescence Microscopy

Wing discs were imaged using an inverted Leica SP8 confocal microscope (Leica Microsystems, Wetzlar, Germany) using the 20x/0.75 NA objective (whole wing disc) or the 63x/1.3 NA (magnified apical sections). Images of the whole wing pouch section depicted in Fig. 2b were acquired using a Leica STELLARIS 5 system (40x/1.25 NA)

### Image processing and analysis

#### Compartment size (wing discs and adult wings)

Z-stacks of the wing discs were projected using “Maximum intensity” algorithm in Fiji and used to measure the area of the posterior compartment (mCherry^+^ signal) and the whole disc (E-cad^+^ silhouette) with the selection tool. Anterior compartment area was calculated by subtracting the two measures in Excel, where the Posterior/Anterior area ratio was also calculated. The compartment ratio in the adult wing was measured using the L4 vein as the compartmental border, which is placed posterior to the actual and comparatively irregular border (Hama et al., 1990), calculating the anterior area again via subtraction from the whole wing surface, before obtaining the Posterior/Anterior area ratio.

#### Cell death

Z-stacks of six slices corresponding to the basal region of the wing discs were projected using the average intensity method in Fiji. mCherry and E-cad antibody signals were used to generate masks of the posterior and whole wing disc pouch using the H/H fold as the dorsal edge. For quantification of apoptosis at the whole pouch (Fig. S1), only the H/H fold detected with E-cad antibody was used. Dcp-1 signal was processed with an automatic image script on MATLAB. Briefly, the image was smoothened using Gauss filtering, and a background image was creating using rolling ball structural elements to perform a partial background subtraction. The resulting image was binarized using automatic threshold, and the signal corresponding to the anterior and posterior compartments of the pouch was isolated using the existing region masks. Percentage of apoptotic area per compartment was obtained using the Regionprops function, and the excess of posterior compartment apoptosis was calculating subtracting the percentages within each compartment.

#### JNK-signalling

Z-stacks of 6 slices spanning the apical region of the wing disc proper at the dorsal region of the pouch were projected using the “average intensity” in Fiji. LacZ intensity was read three times at both the anterior and posterior compartments (the latter identified by the signal of mCherry::hTau) to obtain averages that were used to obtain the Posterior/Anterior ratio for LacZ signal.

### Other processing

Images of the whole discs (e.g., Fig. 1b) were generated with the “maximum projection” algorithm in Fiji. Sagittal projections (Fig. 2b) were made using the “reslice” tool in Fiji, with a maximum projection of 3.78 µm. Images of the dorsal (apoptosis) or apical (JNK-signalling) regions corresponded to representative images of the same averaged projections used for the analyses. Representative images were subjected to minimum modification using Fiji, such as rotation, cropping or automated adjustment of brightness and contrast of the whole view for optimal presentation.

### Statistical analyses

Statistical analyses were performed in Prism 10 (GraphPad). All the datasets were tested for normality using the D’Agostino and Pearson test. Nonparametric tests were used when at least one sample did not display normal distribution, and appropriate corrections were applied if the assumption of equality of SDs was not met. For commonly used tests, such as the t test, two-tailed versions were used. Fig. legends include information about precise *n* numbers, presented data and the type of statistical test (e.g., Mean and standard deviation), with all the individual datapoints presented in the graphs. Quantified differences were visually depicted in the graphs with symbols (*p < 0.05, **p < 0.001, ***p < 0.0001, ****p < 0.00001, ns for no differences between samples).

#### Posterior:anterior compartment size ratio

Posterior:anterior compartment size ratios of larval wing discs and adult wings expressing UAS-driven constructs were compared using the Brown-Forsythe and Welch analysis of variance (ANOVA) test or the Kruskal-Wallis test when normality conditions were not met for at least one of the samples.

#### Apoptotic area

The excess of Dcp-1–positive surface at the posterior compartment was compared between genotypes using the Brown-Forsythe and Welch analysis of variance (ANOVA) test. Additionally, each sample was tested against 0 (no difference in apoptotic area between anterior and posterior compartment) using the one-sample t-test (one-sample Wilcoxon test for samples not following normal distribution).

#### puc-LacZ intensity

Posterior:anterior ratio of β-galactosidase antibody signal was compared between genotypes using the Kruskal-Wallis test.

## Acknowledgements

We thank Natalia Bulgakova for fly reagents. We thank the Imaging and Microscopy Centre (IMC) at Biological Sciences and the technical staff at the invertebrate facility for their support and assistance. We further thank the members of the Mudher lab and the School of Biological Sciences (University of Southampton) for their feedback on the project and manuscript.

## Autor contribution

M.R.M and A.M. designed the project. T.L., J.L, S.A and D.A. generated transgenic flies. M.R.M., E.M.C.S., L.S. and A.M. designed the experimental plan. M.R.M. and A.S.C. conducted the experiments. M.R.M. and A.S.C. analysed the data. M.R.M. and A.M. wrote the manuscript.

## Data availability

The generated *Drosophila* strains are available upon request. Custom automated scripts are available at https://github.com/miguelramirezmoreno/toxicity_assay.

## List of abbreviations

A: Anterior (compartment).
AD: Alzheimer’s Disease.
ADNP: Activity-Dependent Neuroprotective Protein.
*en*: *engrailed* (gene).
FTDP-17: frontotemporal dementia with parkinsonism linked to chromosome 17.
MAP: Microtubule Associated Protein.
mCh: mCherry-tagged (hTau protein).
MTBD: microtubule-binding domain.
NFT: Neurofibrillary Tangles.
P: Posterior (compartment).

## Reagents Table

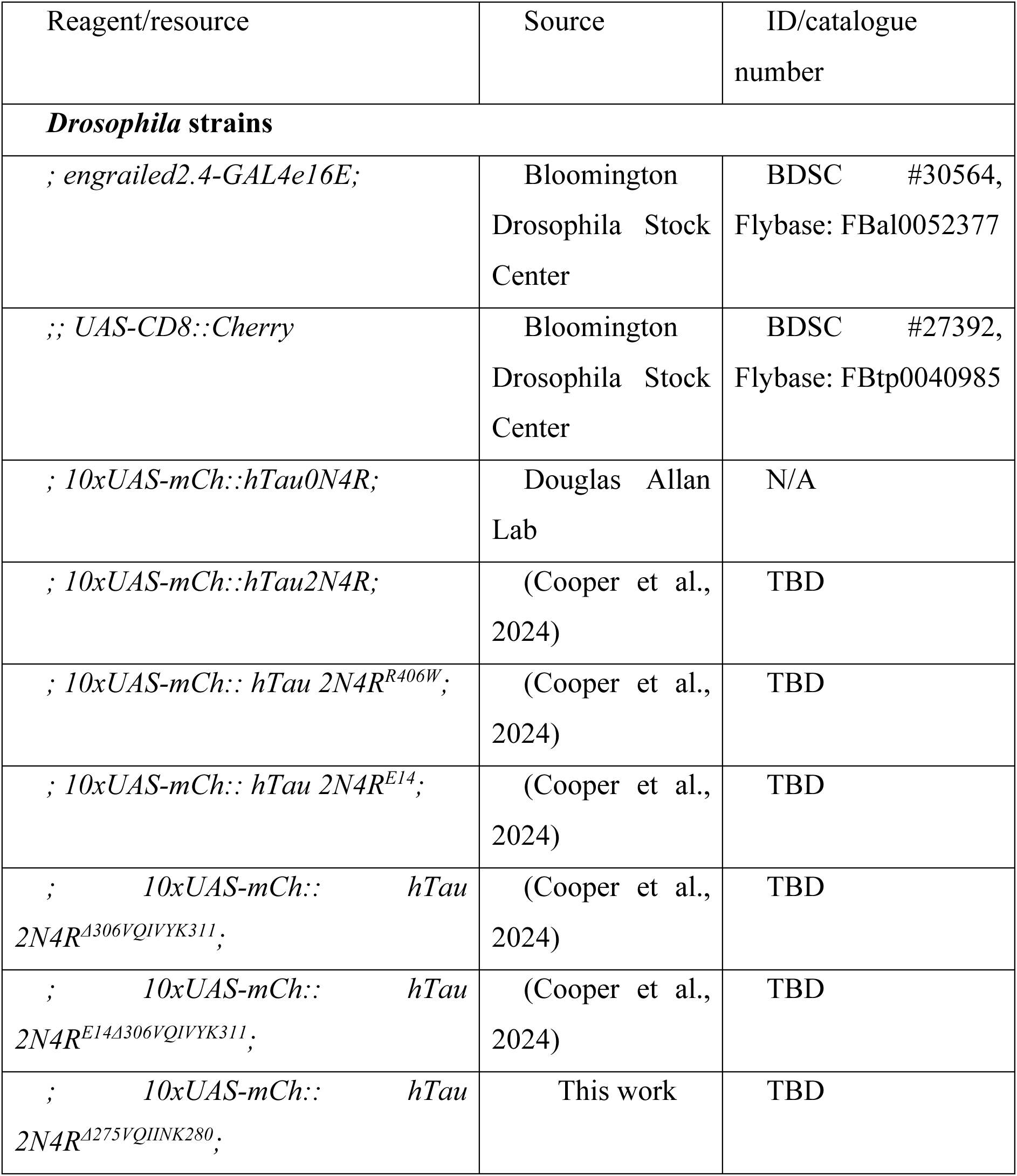
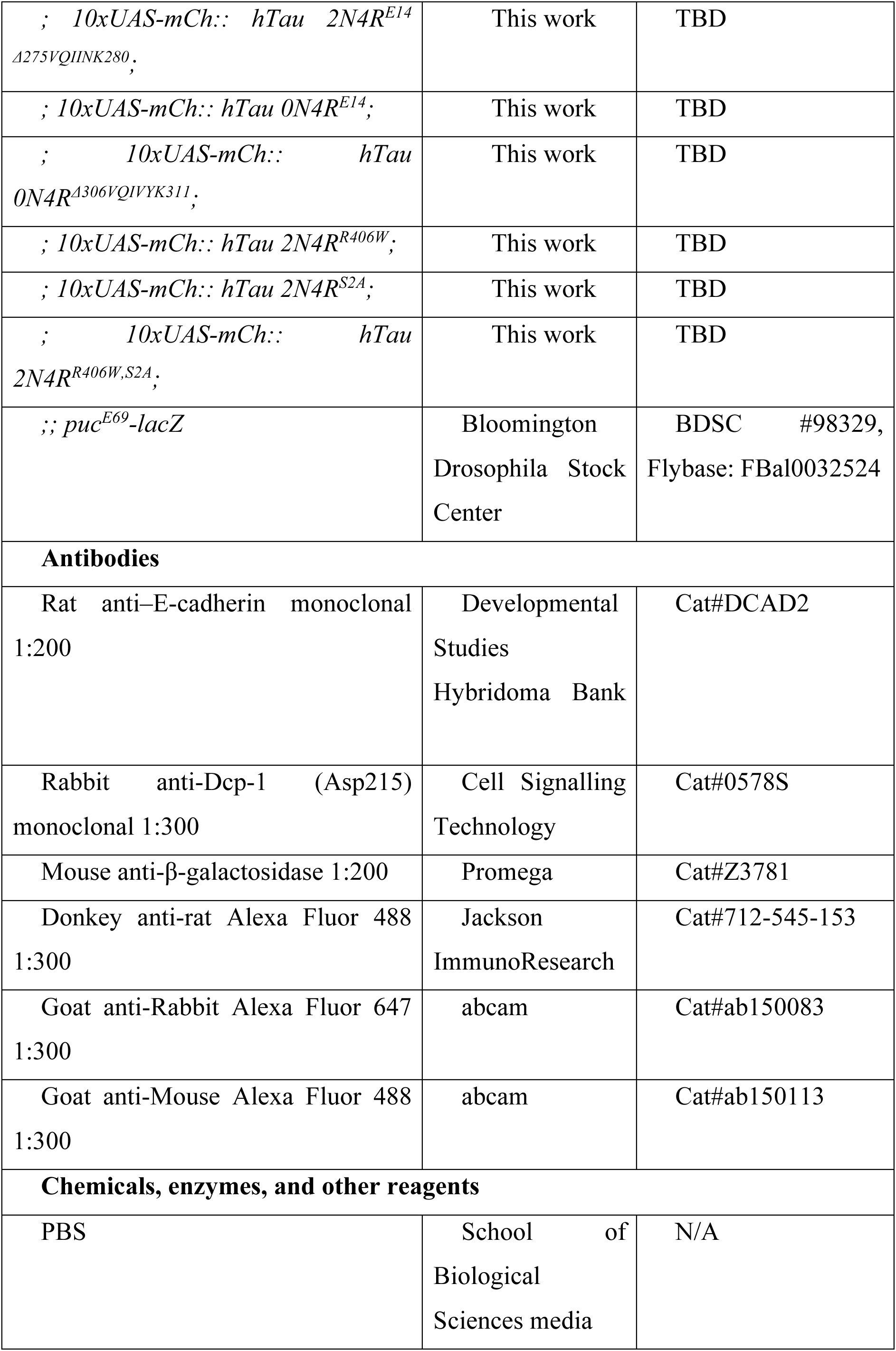
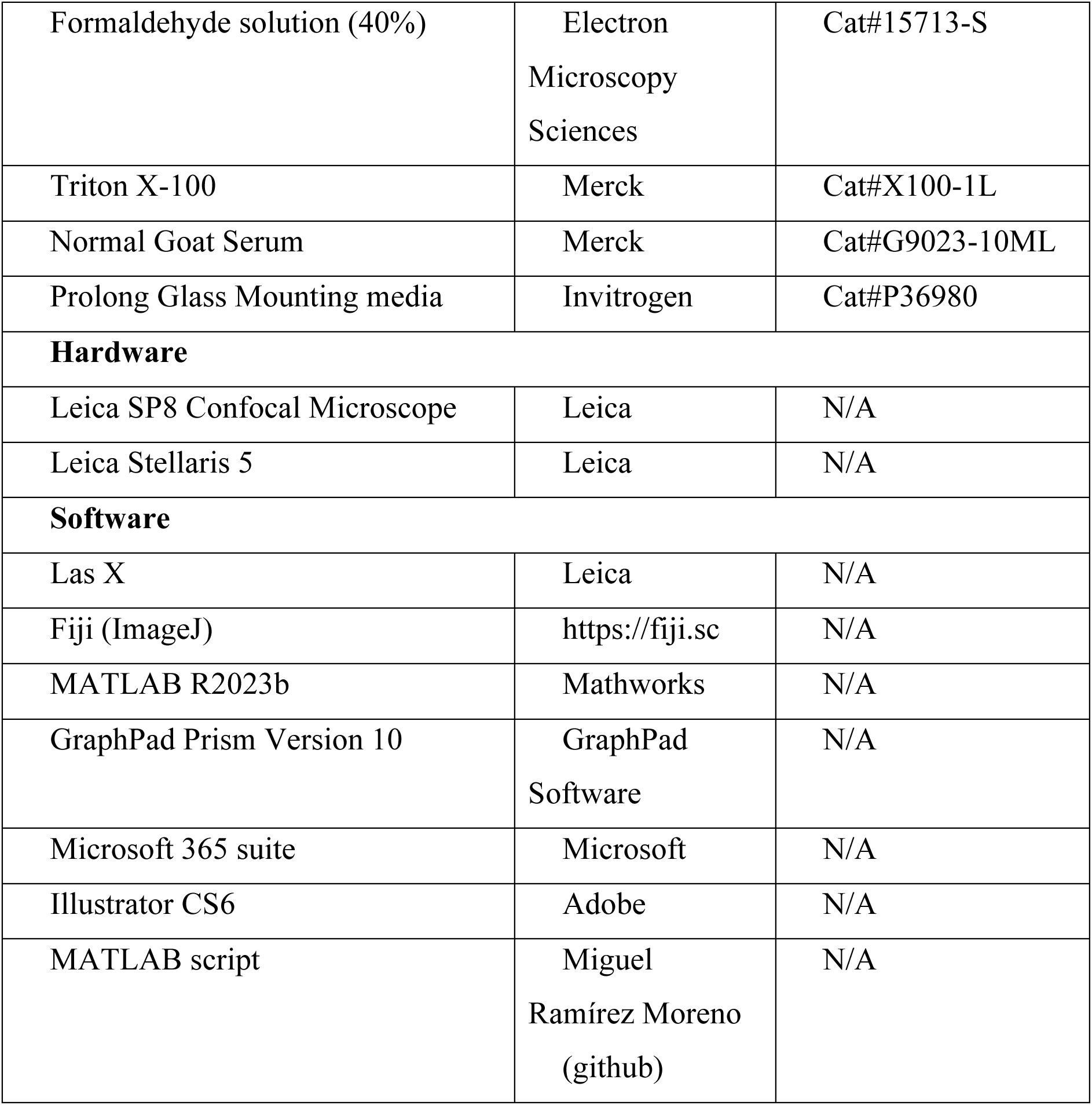

---

**Fig. S1.**
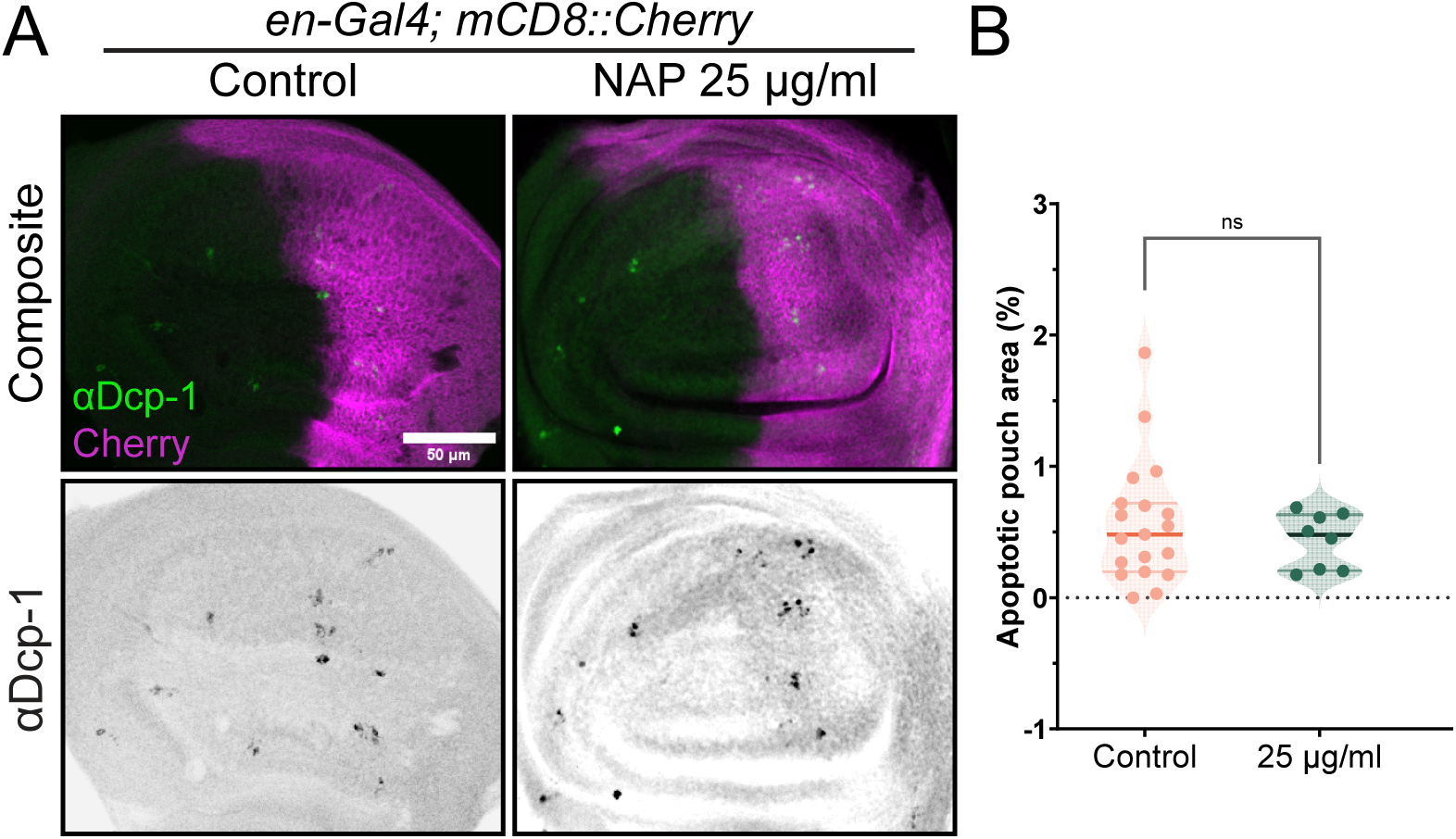
NAP peptide is not toxic for the wing disc cells. (A) Wing pouches expressing mCD8::Cherry at the posterior compartment from larvae raised on normal diet (Control, left) or with NAP peptide (right), showing cleaved Dcp-1 antibody (green, top; grayscale, bottom) and mCherry (magenta, top). Magenta dashed lines at the bottom panel indicate the Anterior-Posterior compartment boundary. Scale bar: 50 µm. (B) Percentage of apoptotic area at the whole wing pouch for the different diet conditions. Dots represent individual discs (n= 19 and 8). Populations are identical according to Mann-Whitney *U* test (p = 0.6964).

**Fig. S2.**
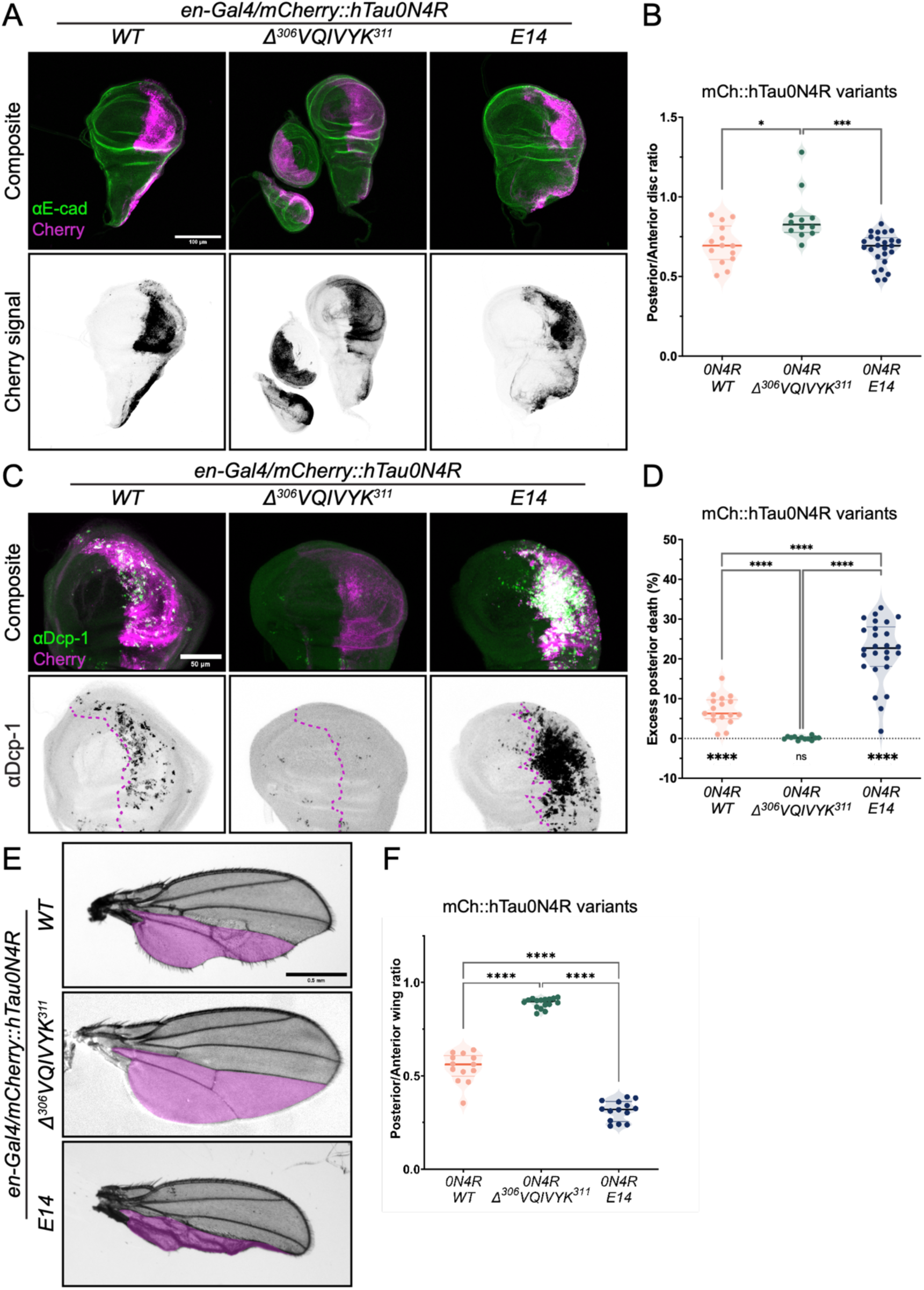
Aggregation and phosphorylation modify 0N4R-induced phenotypes. (A) Wing discs expressing mCherry-tagged (mCh::) variants (columns) of the original hTau0N4R isoform (WT, left), showing mCherry signal (magenta, top; grayscale, bottom) and E-cadherin (E-cad) staining (green, top). Scale bar: 100 µm. (B) Posterior/Anterior ratios of wing discs expressing mCh::hTau0N4R variants, showing design combinations at the bottom of the table. Dots represent individual discs (n= 14, 12 and 25 discs). *p<0.05 and ***p<0.001 (Kruskal-Wallis, test). (C) Wing pouches expressing mCh::hTau0N4R variants, showing cleaved Dcp-1 antibody (green, top; grayscale, bottom) and mCherry (magenta, top) signal. Magenta dashed lines at the bottom panel indicate the Anterior-Posterior compartment boundary. Scale bar: 50 µm. (D) Excess of apoptotic area at the posterior compartment for the indicated genotypes. Dots represent individual discs (n= 17, 12 and 26). ****p<0.0001 (Kruskal-Wallis, and one-sample Wilcoxon test comparison to zero for random distribution of apoptosis). (E) Adult wings (grayscale) expressing mCh::hTau0N4R variants, showing a magenta overlay of the approximated posterior compartment (below L4). Scale bar: 0.5 mm. (F) Posterior/Anterior ratios of adult wings expressing mCh::hTau0N4R variants during development. Dots represent individual wings (n= 13, 16 and 14). ****p<0.001 (Kruskal-Wallis test).

